# Toward Precision Molecular Surgery: Robust, Selective Induction of Microhomology-mediated End Joining *in vivo*

**DOI:** 10.1101/291187

**Authors:** Hirotaka Ata, Thomas L. Ekstrom, Gabriel Martínez-Gálvez, Carla M. Mann, Alexey V. Dvornikov, Kyle J. Schaefbauer, Alvin C. Ma, Drena Dobbs, Karl J. Clark, Stephen C. Ekker

**Affiliations:** Mayo Clinic, Center for Clinical and Translational Science. Rochester, MN, USA; Mayo Clinic, Graduate School of Biomedical Sciences. Rochester, MN, USA; Mayo Clinic, Medical Scientist Training Program, Rochester, MN, USA; Iowa State/Mayo Clinic Alliance for Genome Engineering; Iowa State University, Genetics, Development, and Cell Biology, Ames, IA, USA; Mayo Clinic, Biochemistry and Molecular Biology. Rochester, MN, USA; The University of Hong Kong, Department of Paediatrics and Adolescent Medicine, Hong Kong, Hong Kong

## Abstract

One key problem in precision genome editing is the resultant unpredictable plurality of sequence outcomes at the site of targeted DNA double-strand breaks (DSBs). This is due to the typical activation of the versatile Non-homologous End Joining (NHEJ) pathway. Such unpredictability limits the utility of somatic gene editing for applications including gene therapy and functional genomics. For germline editing work, the accurate reproduction of identical alleles using NHEJ is a labor intensive process. In this study, we propose inducing Microhomology-mediated End Joining (MMEJ) as a viable solution for improving somatic sequence homogeneity *in vivo*, capable of generating a single predictable allele at high rates (56% ~ 86% of the entire mutant allele pool). Using a combined dataset from zebrafish (*Danio rerio*) *in vivo* and human HeLa cell *in vitro* as a training dataset, we identified specific contextual sequence determinants surrounding genomic DSBs for robust MMEJ pathway activation. We then applied our observation and prospectively designed MMEJ-inducing sgRNAs against a variety of proof-of-principle genes and demonstrated a high level of mutant allele homogeneity at these loci. F0 mutant zebrafish embryos and larvae generated with these gRNAs faithfully recapitulated previously reported, recessive loss-of-function phenotypes. We also provide a novel algorithm MENTHU (http://genesculpt.org/menthu/) for improved prediction of candidate MMEJ loci, suitable for both targeted and genome-wide applications. We believe that this MMEJ-centric approach will have a broad impact on genome engineering and its applications. For example, whereas somatic mosaicism hinders efficient recreation of a knockout mutant allele at base pair resolution via the standard NHEJ-based approach, we demonstrate that F0 founders transmitted the identical MMEJ allele of interest at high rates. Most importantly, the ability to directly dictate the reading frame of an endogenous target will have important implications for gene therapy applications in human genetic diseases.

**Author Summary:** New gene editing tools precisely break DNA at pre-defined genomic locations, but cells repair these lesions using diverse pathways that often lead to unpredictable outcomes in the resulting DNA sequences. This sequence diversity in gene editing outcomes represents an important obstacle to the application of this technology for human therapies. Using a vertebrate animal as a model system, we provide strong evidence that we can overcome this obstacle by selectively directing DNA repair of double-stranded breaks through a lesser-described pathway termed Microhomology-mediated End Joining (MMEJ). Unlike other, better-understood pathways, MMEJ uses recurring short sequence patterns surrounding the site of DNA breakage. This enables the prediction of repair outcomes with improved accuracy. Importantly, we also show that preferential activation of MMEJ is compatible with effective gene editing. Finally, we provide a simple algorithm and software for designing DNA-breaking reagents that have high chance of activating the MMEJ pathway. We believe that the MMEJ-centric approach to be broadly applicable for a variety of gene editing applications both within the laboratory and for human therapies.

**Author Contribution:** **HA** contributed in Conceptualization, Data Curation, Formal Analysis, Investigation, Funding Acquisition, Methodology, Validation, Visualization, Writing – Original draft preparation, and Writing – Review and Editing. **TLE** contributed in Data Curation, Investigation, Writing – Original draft preparation, and Writing – Review and Editing. **GMG** contributed in Software, Validation, and Writing. **CMM** contributed in Software Validation, and Writing. **AVD** contributed in Investigation, Methodology, Validation, and Writing – Review and Editing. **KJS** contributed in Investigation and Writing – Review and Editing. **ACM** contributed in Conceptualization, Data Curation, Investigation, and Writing – Review and Editing. **DD** contributed in Funding Acquisition, Resources, and Writing – Review and Editing. **KJC** contributed in Conceptualization, Funding Acquisition, Resources, Supervision, and Writing – Review and Editing. **SCE** contributed in Conceptualization, Funding Acquisition, Project Administration, Resources, Supervision, Writing – Review and Editing.

## Introduction

Programmable nucleases such as TALEN (Transcription Activator-like Effector Nuclease) and CRISPR (Clustered Regularly Interspaced Short Palindromic Repeats) systems have enabled a new era of scientific research(1, 2). Instead of relying on knock-down models or expensively outsourced knockout lines, laboratories across the world now have tools with which to generate indels (Insertions and deletions) of varying sizes in the gene(s) of interest. However, DNA Double-strand Break (DSB) repairs largely result in diverse sequence outcomes owing to the unpredictable nature of the most commonly used Non-homologous End Joining (NHEJ) pathway(3, 4) (**Fig 1**). This significantly confounds experimental readouts because knock-out cell lines often harbor more than just one desired frameshift mutation. In the case of model organisms such as zebrafish (*Danio rerio*), the F0 founders are genetically mosaic, warranting a complex and time-consuming series of outcrossing to establish molecularly defined lines before any biological questions can be addressed(5, 6).

**Fig 1.**
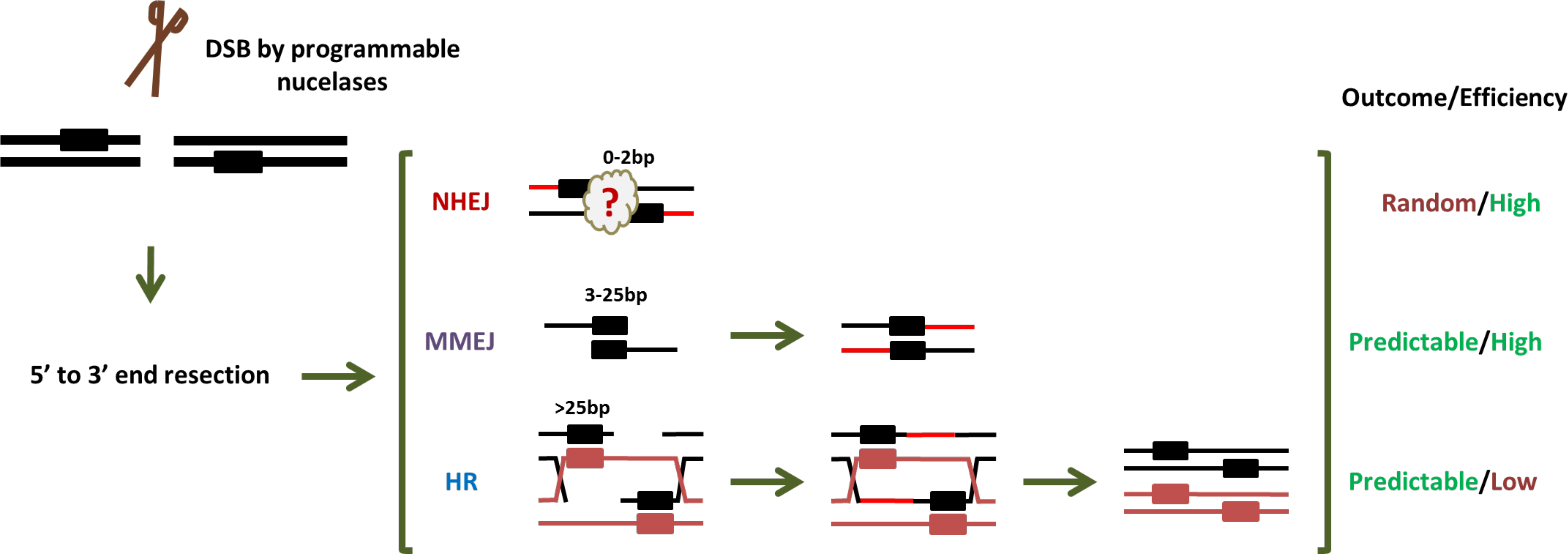
MMEJ is a unique DSB repair pathway that results in highly efficient and highly stereotyped mutagenesis. DSB repair generally begins with end resections to varying degrees. Once the ends are processed, end joining in NHEJ often occurs between incompatible ends, producing unpredictable and heterogeneous mutant alleles. In contrast, MMEJ uses region of sequence microhomology flanking a DSB to temporarily appose the two strands. A polymerase will then elongate DNA from the homology arms in a templated fashion, resulting in predictable mutagenesis. Although both MMEJ and NHEJ can lead to high efficiency, biallelic mutagenesis of a target gene, HDR is usually a low-efficiency, monoallelic process. This is because DSB is repaired by recruiting homologous DNA as a template for repair. Of the HDR pathways, HR results in a high-fidelity repair, usually confined to late S ~ G2 phase because of the proximity requirement of the two chromosomes. Rectangular boxes represent homology arms of varying lengths indicated above them.

In contrast to NHEJ, the MMEJ (Microhomology-mediated End Joining) DNA repair pathway utilizes a pair of locally available direct sequence repeats on both sides of a DSB that are apposed, annealed and extended(7-10) (**Fig 1**). As such, DSB repair outcomes are highly stereotyped, resulting in deletion of the intervening sequence as well as one of the repeats. Consequentially, there is an increasing interest in utilizing MMEJ for precision genome engineering applications(11-14). To date, however, effective harnessing of this pathway remains challenging due to the paucity of genetic and mechanistic understanding(8).

Bae *et al*.(14) developed a sequence-based scoring system to estimate the frequency of MMEJ-associated deletions induced by DSBs in human cells. While this improved the predictability of MMEJ activation, the DSB repair outcomes tended to consist of a heterogeneous population of multiple MMEJ alleles. In this study, we sought to improve upon the existing algorithm with the goal of developing tools to more reliably predict target loci that would be predisposed to generate a more homogeneous mutant allele population through MMEJ. We demonstrate the feasibility and utility of such reagent design on the molecular level (i.e., DNA repair outcomes) and on the physiological level (i.e., F0 phenotype). We believe our approach can inform and benefit applications such as rapid phenotype-genotype correlation in F0 animals, with an eye toward applications in human gene therapy and facilitation of resource sharing and recreation of various cell and animal lines on a global scale.

## Results

### MMEJ is an Active Repair Pathway in the Genetically Unaltered Zebrafish Embryo

Prior work examining MMEJ activation in vertebrate organisms primarily focused on *in vitro* models (8-10, 14-18). Initial analyses using a targeted knockin strategy suggested that MMEJ was operational in the zebrafish embryo, though the efficiency of these MMEJ outcomes was rather modest(13). One study reported incidental identification of MMEJ inductions at two zebrafish genomic loci using programmable nucleases (19). However, no consortium – small or large – of genomic loci that repair primarily through NHEJ vs MMEJ has been compiled. To this end, we examined the repair outcomes of previously designed TALEN and CRISPR-Cas9 genomic reagents (**S1 Table**). The plurality of custom enzymes induced diverse sequence outcomes, consistent with NHEJ being used as the primary DNA repair pathway. However, a few reagents induced sequence outcomes satisfying the following criteria, suggesting that MMEJ was the preferred repair pathway: 1) most predominant allele is the top predicted allele by the Bae *et al*. algorithm(14); 2) most predominant allele comprises ≥ 50 % of the total mutant allele population; and 3) mutagenic efficiency > 20%. For the purpose of this study, a programmable nuclease satisfying all these criteria is referred to as a “Winner-Take-All” reagent. Three TALEN (*chrd*, *mitfa* #4 & *surf1*) and two CRISPR-Cas9 (*surf1* & *tyr* #2) reagents fell into this category (**S1 Table**, **Fig 2A**, **Fig 3A**).

**Fig 2.**
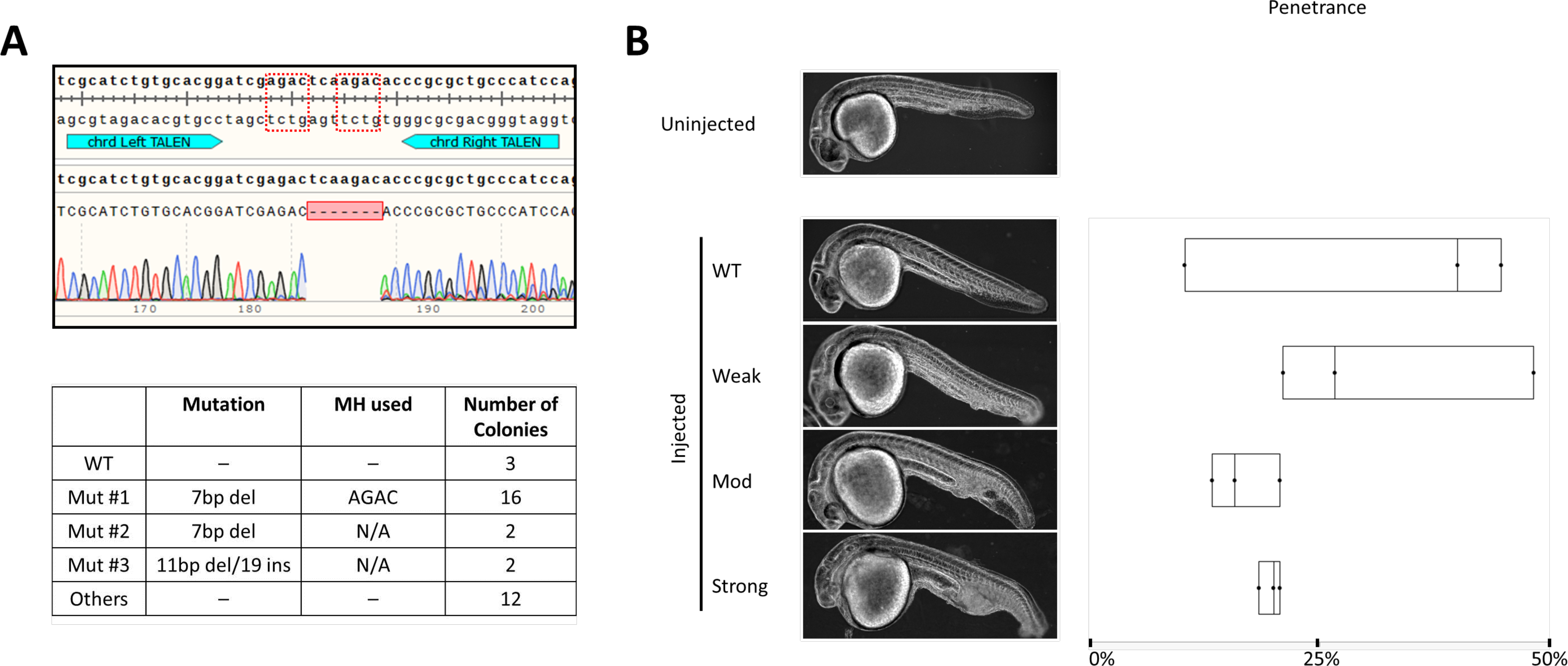
Winner-Take-All TALEN reagent can be used to recapitulate previously reported loss-of-*chrd*-function phenotype in 1 dpf F0, injected larvae. **A** *Top* – Wildtype *chrd* sequence with TALEN binding sites annotated in teal. The dotted red boxes are MH arms predicted to be used most frequently. Raw sequence alignment of the whole PCR amplicon demonstrates that the majority of reads are the expected 7bp deletion allele. *Bottom* – summary data from subcloning analyses. 50% of the mutant allele recovered were of the predicted MH allele. **B** Previously reported *chrd* loss-of-function phenotype was successfully recapitulated using this TALEN pair. Phenotype severity was graded by the degree of ICM expansion in the tail and by the reduced head size by 1 dpf. Box plot demonstrating phenotypic penetrance is provided. N = 3 biological and technical replicates. At least 29 injected animals were scored in each experiment.

**Fig 3.**
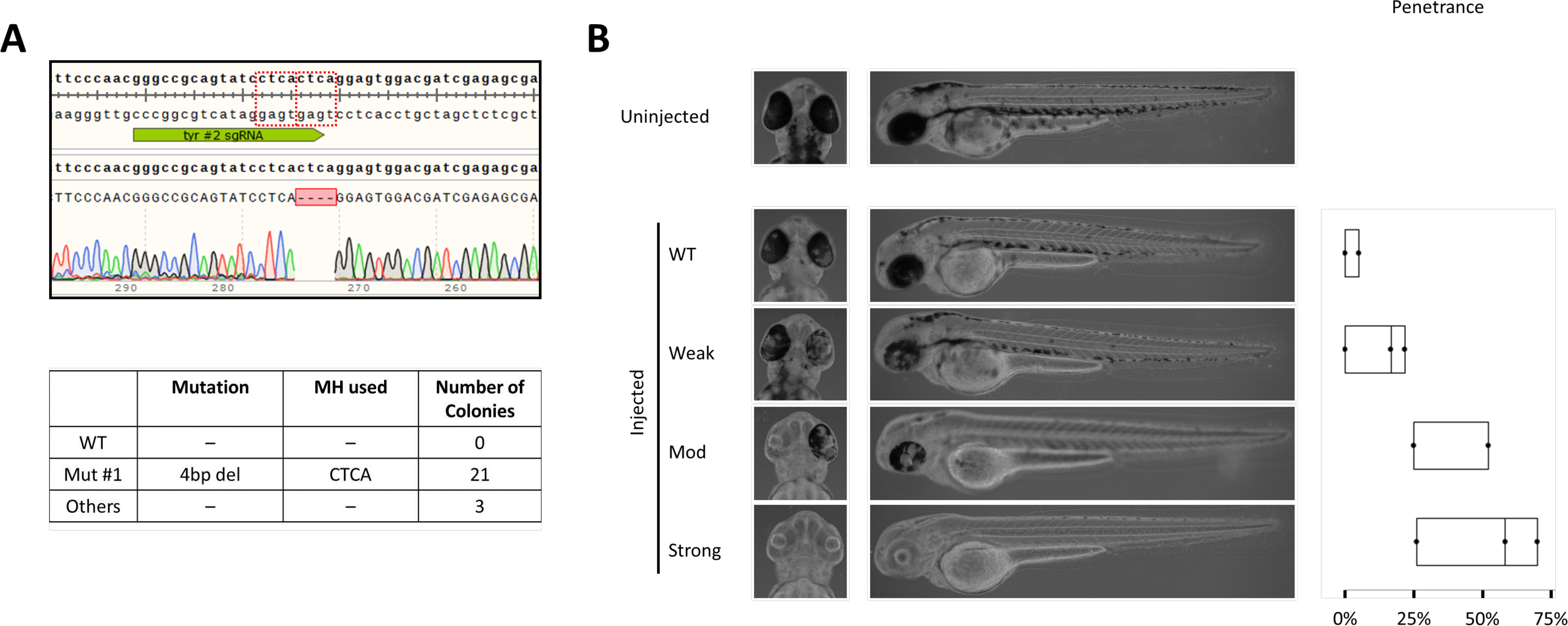
Winner-Take-All sgRNA against *tyr* can be used to recapitulate loss-of-melanophore phenotype in 2 dpf, injected F0 larvae injected. **A** *Top* – Wildtype *tyr* sequence with the #2 sgRNA target site annotated in green. The dotted red boxes are MH arms predicted to be used most frequently. Raw sequence alignment of the whole PCR amplicon demonstrates that the majority of reads are the expected 4bp deletion allele. *Bottom* – summary data from subcloning analyses. 88% of the mutant allele recovered were of the predicted MH allele. **B** Previously reported *tyr* loss-of-function phenotype was successfully recapitulated using this CRISPR-Cas9. Phenotype severity was graded by the loss of retinal pigmentation. Partial loss of retinal pigmentation was considered a Weak phenotype, whereas complete loss of pigmentation in one or both eyes were considered Moderate and Strong phenotypes, respectively. Box plot demonstrating phenotypic penetrance is provided. N = 3 biological and technical replicates. At least 12 injected animals were scored in each experiment.

Injecting the *chrd* TALEN pair (37.5 pg/arm) resulted in characteristic *chrd* loss of function phenotypes: Intermediate-Cell-Mass (ICM) expansion and a smaller head by 1 day post-fertilization(20) (1 dpf; **Fig 2B**). Median penetrance for Moderate and Severe phenotypes was 15.8% and 20.0%, respectively (**Fig 2B**, **S2 Table**). Strong MMEJ activation by this TALEN pair was confirmed by subcloning results (**Fig 2A**) – 16/32 recovered mutant reads corresponded to the top predicted 7bp deletion allele. Similarly, perturbing *tyr* gene with a CRISPR-Cas9 reagent recapitulated a previously reported loss of melanin production phenotype, observable by 2 dpf(21) (**Fig 3B**). Ribonucleoprotein (RNP) delivery at the dose of 300 pg *tyr* #2 sgRNA and 660 pg Cas9 resulted in Moderate and Severe loss of pigmentation phenotypes in 22.7% and 50.0% of embryos, respectively (**Fig 3B**, **S2 Table**). Subcloning analysis showed 21/24 (88%; **Fig 3A**) of resulting alleles contained a 4bp deletion consistent, with strong MMEJ activation by this CRISPR-Cas9. Together with the *chrd* TALEN results, these data support that MMEJ can be an effective repair pathway in F0 embryos at some genomic loci.

### Many Bae *et al*. Predicted MMEJ Loci Are Preferentially Repaired by NHEJ

A subset of the zebrafish reagents described above was prospectively designed using the Bae *et al*. algorithm(14) (**S1 Table**). This algorithm calculates the strength of each pair of microhomology arms according to the length and GC content of each pair, as well as the length of the intervening sequence (i.e., *Pattern Score*). The additive sum of all the possible *Pattern Scores* is then returned as *Microhomology Score*. This latter score was found to have positive correlation with the rate of MMEJ activation in HeLa cells(14). All fourteen prospectively designed reagents had a *Microhomology Score* of at least 4000 – a median score found on human *BRCA1* gene. However, only four of these reagents induced majority MMEJ outcomes as judged by the *Microhomology Fraction* (**S1 Table**, **S1 Note**). We therefore retrospectively analyzed the repair outcomes of these reagents to identify additional factor(s) that could enhance predictability of MMEJ induction.

### Rate of Pattern Score Change as a Discrimination Factor for MMEJ Induction *in vivo* and *in vitro*

Intriguingly, when the pattern score values clustered closely to one another (i.e., a flatter *Slope Value* as calculated according to **S2 Note**), this was indicative of an unfavorable target for MMEJ activation in zebrafish embryos. Conversely, loci at which pattern scores dropped precipitously (i.e., a steeper *Slope Value*) were good candidates for MMEJ activation *in vivo* (p = 0.0048; **S1 Figure**). Based on these observations, we hypothesized that locally available microhomology pairs are in direct competition with one another, such that overabundance of these pairs is a negative predictor of MMEJ activation. In other words, MMEJ activation is more favorable at loci with only one or two predominant microhomology pair(s) (Low Competition loci) rather than many strong microhomology pairs (High Competition loci).

To determine whether the zebrafish-based hypothesis was generalizable to human cells (HeLa), we re-analyzed the deep sequencing dataset used to generate the Bae *et a.l* algorithm(14). Available results from 90 genomic loci were sorted alphabetically by the names of target genes. Outcomes from the first 50 targets showed a correlation similar to that observed in zebrafish; higher *Microhomology Fractions* generally correlated with low *Slope Values* from the first 50, alphabetically sorted targets (p = 0.00001; **S2 Figure A**). This correlation was lost when microhomology arms of 2 bp were included in the analysis (p = 0.2644; **S2 Figure B**). Accordingly, microhomology arms of less than 3 bp were excluded from subsequent analyses. The remaining 40 targets were then binned into High, Medium and Low Competition groups based on quartiles (**S2 Figure C**) – the median *Microhomology Fraction* was significantly higher in the High Competition group than in the Low Competition group (0.300 vs 0.105, p = 0.011; **S2 Figure D**).

### Competition Hypothesis Predicts New Winner-Takes-All Reagents

Based on this Competition Hypothesis, we designed and analyzed the DSB repair outcomes of 20 Low Competition sgRNA targets across 9 genes (**S3 Table**). *Slope Values* smaller than −40 was used as the cut-off for Low Competition, as 3 out of 4 previously designed zebrafish targets produced majority MMEJ outcomes in this range (**S1 Table and S1 Figure**). For initial assessments, we used TIDE (Tracking Indels by DEcomposition) analysis – a chromatogram analyzing tool that estimates proportions of length varying mutant alleles present in a pool of mixed alleles (22) – which revealed that 5 of these sgRNAs against 3 genes: *mtg1*, Mitochondrial GTPase 1; *tdgf1*, Teratocarcinoma-Derived Growth Factor 1; and *ttn.2*, titin (*ttn.2 #1*, *ttn.2 #2*, *ttn.2* N2B #1) were in the “Winner-Take-All” class. These results were subsequently confirmed by subcloning analysis (**S3 Table**). Perturbation of *tdgf1* (alternatively known as *One-eyed Pinhead*) causes aberrant “fused eyes” morphology and cyclopia, as judged by reduced forebrain protrusion by 1 dpf(23) (**Fig 4B**). Aberrant head morphology alone was classified as Weak, whereas that in combination with varying degrees of forebrain protrusion was classified as Moderate or Strong phenotypes. RNP injections of CRISPR-Cas9 at the dose of 300 pg sgRNA and 660 pg Cas9 resulted in median penetrance for Moderate and Severe morphology at 21.8% and 11.4% (**Fig 4B**, **S2 Table**). The subcloning results were consistent with the noted phenotypic outcomes from this common 4bp deletion (**Fig 4A**).

**Fig 4.**
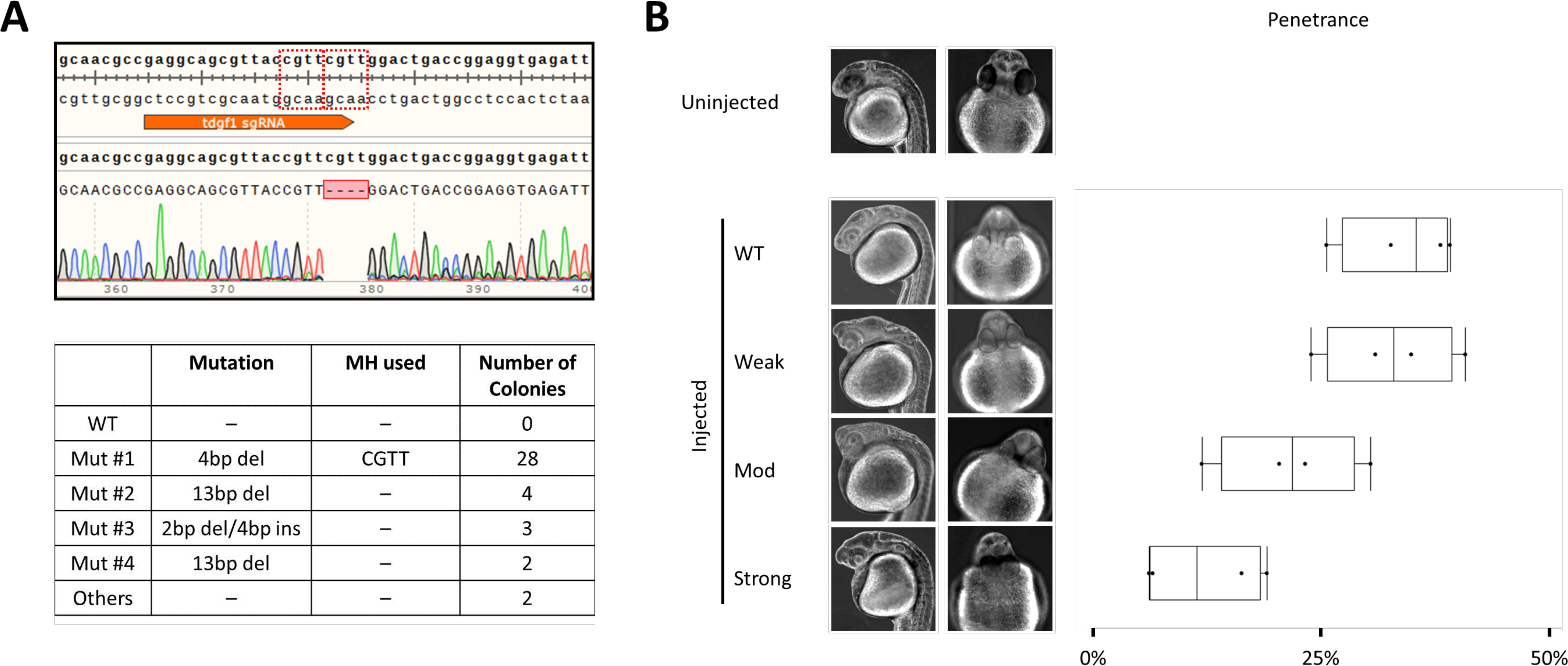
Prospectively designed Winner-Take-All reagent against *tdgf1* can be used to reproduce gross developmental defect in 1 dpf, injected F0 larvae. **A** *Top* – Wildtype *tdgf1* sequence with sgRNA target site annotated in orange. The dotted red boxes are MH arms predicted to be used most frequently. Raw sequence alignment of the whole PCR amplicon demonstrates that the majority of reads are the expected 4bp deletion allele. *Bottom* – summary data from subcloning analyses. 72% of the mutant allele recovered were of the predicted MH allele. **B** Previously reported *tdgf1* loss-of-function phenotype was successfully recapitulated using this CRISPR-Cas9. Phenotype severity was graded by the “pinhead” morphology and cyclopia. Pinhead morphology alone was classified as Weak, whereas Moderate and Strong phenotypes also presented with varying degrees of cyclopia judged by the distance of forebrain protrusion. In the Strong class, the forebrain does not separate the eyes, and they are fused together. Box plot demonstrating phenotypic penetrance is provided. N = 4 with 3 biological and 4 technical replicates. At least 42 injected animals were scored in each experiment.

We next explored whether these “Winner-Take-All” reagents could be useful for recapitulating a more subtle phenotype than the aberrant gross morphologies observed in the *tdgf1* mutants. Splice blockade at the N2B exon of *ttn.2* gene by a synthetic morpholino oligonucleotide was previously reported to reduce cardiac contractility by ~70% on 2 dpf(24), phenocopying the *pickwick*^m171^ mutation(25). RNP delivery at the dose of 300 pg *ttn.2* N2B #2 sgRNA + 660 pg Cas9 resulted in reduction of the shortening fraction to a comparable degree (**Fig 5B**). Importantly, injecting RNP with *tyr #2* sgRNA or sgRNA and Cas9 independently, at the same doses, did not affect the shortening fraction. Due to the high editing efficiency (**Fig 5A**), animals injected with these doses of CRISPR-Cas9 were not viable in post larval phases. For this reason, animals injected at the lower dose of 75 pg sgRNA + 165 pg Cas9 protein were raised to adulthood. Two F0 founders were successfully outcrossed to wild type zebrafish. Heterozygous offspring were identified using the dsDNA heteroduplex-cleaving Surveyor assay(26), and the transmission of the top predicted 5bp deletion allele was confirmed from both founders by subcloning analyses (**S3 Figure**).

**Fig 5.**
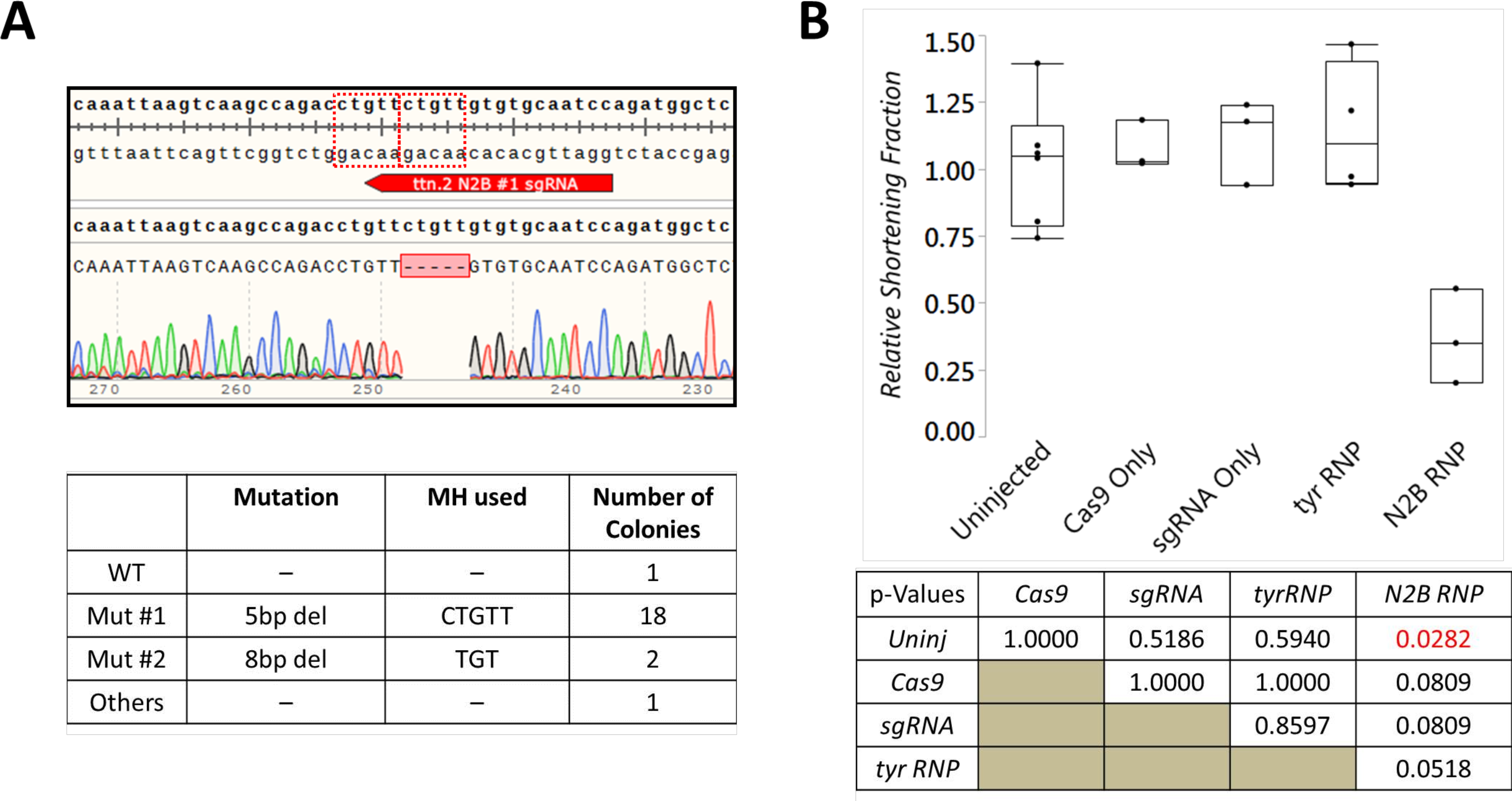
Winner-Take-All reagent against *ttn.2* N2B results in specific reduction of shortening fraction in 2 dpf F0 zebrafish. **A** *Top* – Wildtype *ttn.2* sequence at the N2B exon with sgRNA target site annotated in red. The dotted red boxes are MH arms predicted to be used most frequently. Raw sequence alignment of the whole PCR amplicon demonstrates that the majority of reads are the expected 5bp deletion allele. *Bottom* – summary data from subcloning analyses. 86% of the mutant allele recovered were of the predicted MH allele. **B** Previously reported *pickwick* phenotype was successfully recapitulated using this CRISPR-Cas9. 2 dpf zebrafish were immobilized in 3% methylcellulose for live recording of cardiac functions. Whereas injections with Cas9 only (660pg), N2B #1 sgRNA only (300pg), or *tyr* #2 sgRNA RNP (300pg sgRNA + 660pg Cas9) did not result in changes in shortening fraction at this age, RNP injection containing N2B #1 sgRNA (300pg sgRNA + 660pg Cas9) resulted in a specific reduction in shortening fraction by 65%. N > 3 biological and technical replicates. At least 5 injected animals were scored in each experiment. P-values calculated using the Wilcoxon Each Pair Calculation (adjusted for multiple comparisons)

We also designed an sgRNA against exon 13 of *ttn.2* (*ttn.2* #2 sgRNA), expected to produce a 12 bp deletion allele as a proof-of-principle for in-frame gene correction (**Fig 6A**). RNP delivery at the dose of 300 pg sgRNA + 660 pg Cas9 resulted in the induction of this 12 bp deletion allele at 72.7% of the resultant clones. While the injected animals presented with mild cardiac edema evident by 2 dpf (median rate: 50.0%; **Fig 6B**, **S2 Table**), unlike the N2B sgRNA #1 CRISPR-Cas9 injected animals, these were viable to adult age.

**Fig 6.**
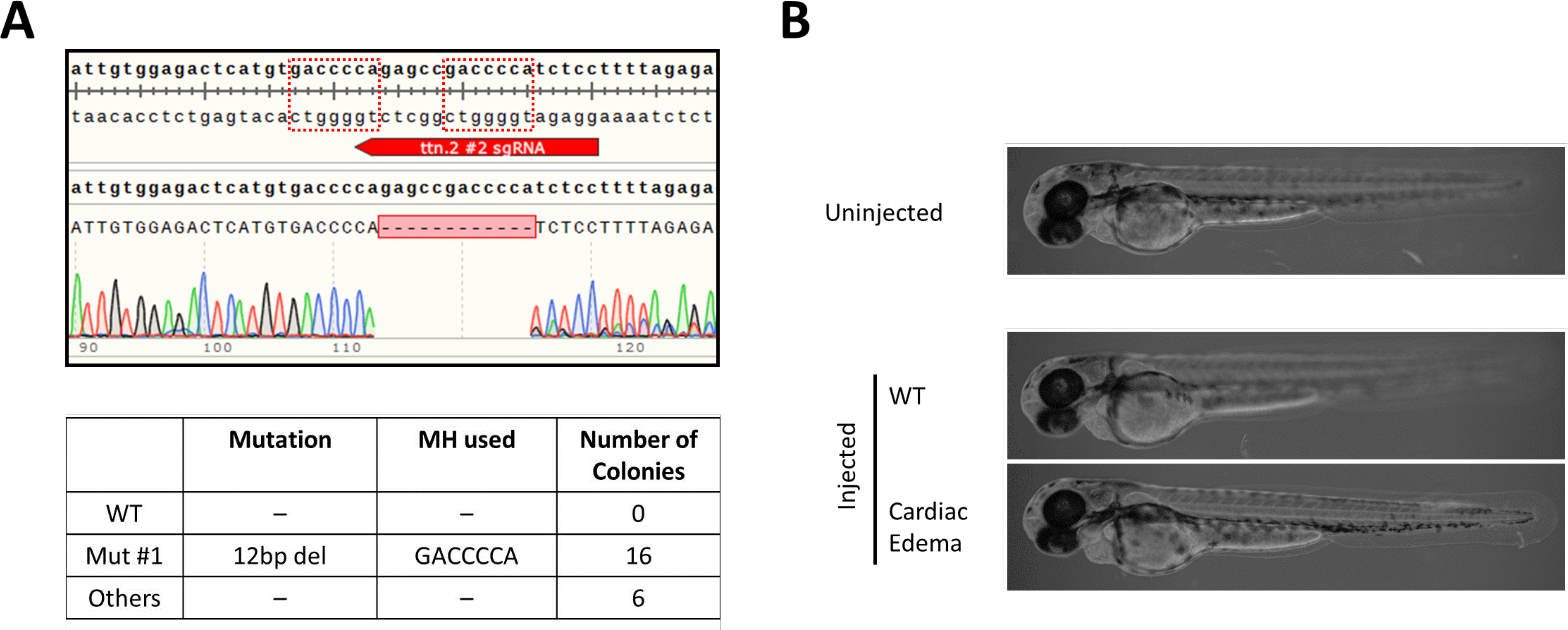
Winner-Take-All reagent can be used for in-frame gene alteration. **A** *Top* – Wildtype *ttn.2* sequence with sgRNA target site annotated in red. The dotted red boxes are MH arms predicted to be used most frequently. Raw sequence alignment of the whole PCR amplicon demonstrates that the majority of reads are the expected 12bp deletion allele. *Bottom* – summary data from subcloning analyses. 73% of the mutant allele recovered were of the predicted MH allele. **B** 2 dpf zebrafish larvae injected with *ttn.2 #2* sgRNA RNP (300pg sgRNA + 660pg Cas9) grossly appear normal with the exception of mild cardiac edema. Median penetrance was 50%. N = 3 biological and technical replicates. At least 9 injected animals were scored in each experiment.

### Low Competition Plus Proximity of Microhomology Arms Strongly Predicts Winner-Take-All Reagents: V2

These data implicate the utility of “Winner-Take-All” class reagents for various applications that require precision gene editing. However, sgRNA design based on the Competition Hypothesis yielded only 5 Winner-Take-All reagents out of 20 that were tested (**S3 Table**, **S3 Note**). Although this represents an improvement over the initial approach relying solely on the *Microhomology Score*, we sought to further fine-tune the predictability for the Winner-Take-All targets. To this end, we pooled the results from all the programmable nucleases described above (**S1 and S3 Tables**) and seven Medium ~ High Competition sgRNAs designed as controls based on the Competition Hypothesis (**S4 Table**). In so doing, we noted that Winner-Take-All outcomes were observed only if the two arms of the microhomology of the top predicted MMEJ allele for a locus were separated by no more than 5 bp of intervening sequence. Thus, we identified a second parameter: high ratio (≥ 1.5) of the *Pattern Scores* between the top and second predicted MMEJ alleles for a given locus (**Fig 7**). Seven out of eight reagents that satisfied both of these parameters were Winner-Take-All. Of the nine reagents that satisfied the first parameter but not the second, 2 were Winner-Take-All. All the other 30 reagents that failed to meet the first parameter failed to induce the top predicted MMEJ allele strongly. Most importantly, all the failed cases, i.e., incorrect predictions according to the original Competition Hypothesis, can be explained using our revised approach (Competition Hypothesis V2; **Fig 7C**). Version 2 also captured 3 Winner-Take-All reagents that would have been missed by the original Competition Hypothesis alone, and 1 Winner-Take-All reagent that would have been missed by the *Microhomology Score* alone. Finally, a similar trend was observed with the HeLa cell dataset (**S4 Figure**). While the effect is not as dramatic as in zebrafish, this points to a possibly conserved mechanism for MMEJ activation in vertebrate organisms.

**Fig 7.**
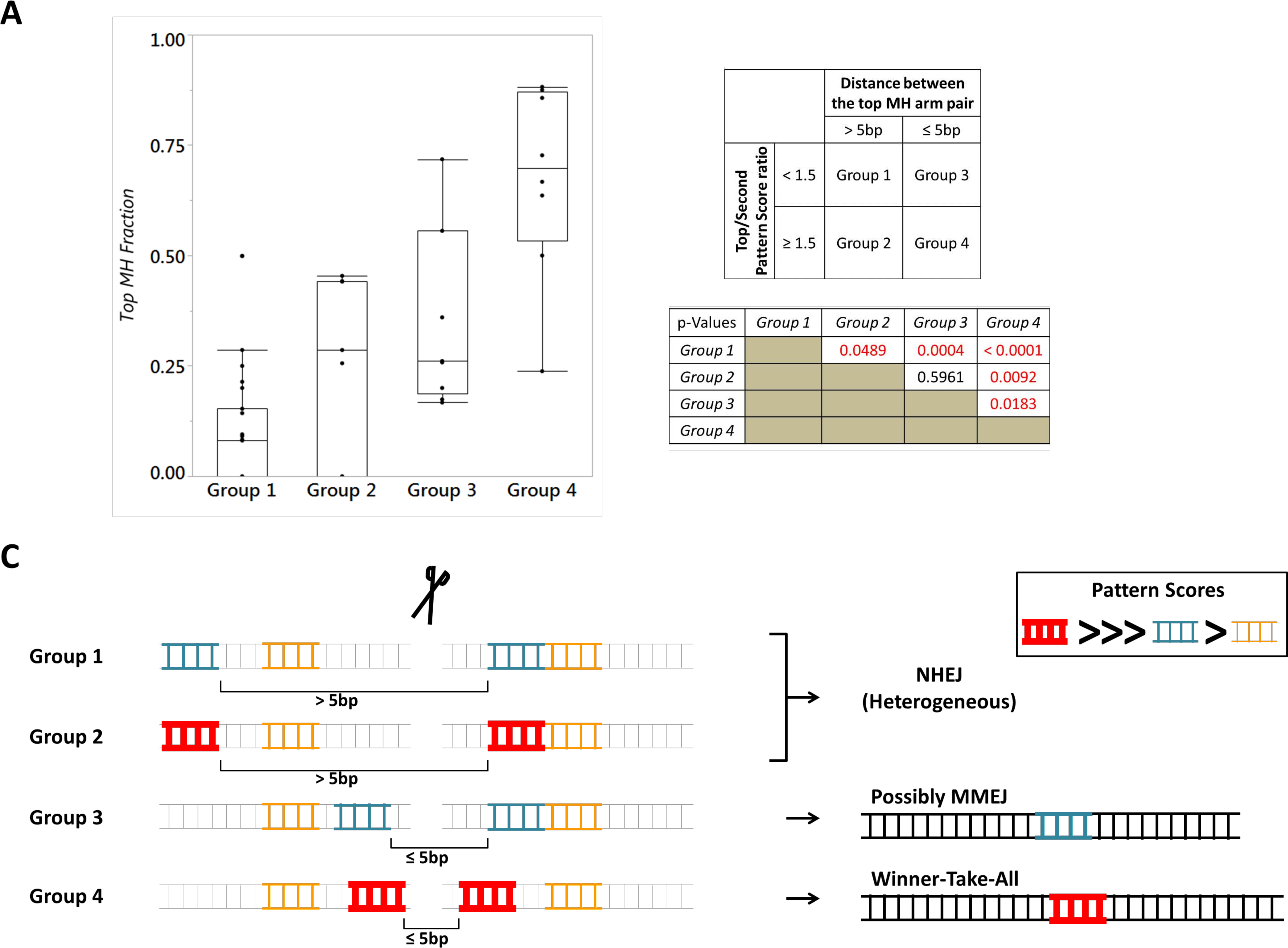
Competition Hypothesis Version 2. **A** Outlier plot summarizing repair outcomes from 47 genomic targets using TALEN and CRISPR-Cas9. Close proximity of 2 MH arms (Groups 3 and 4) appears to be the primary determinant for generating Winner-Take-All type outcomes as no target from Groups 1 and 2 had Top MH Fraction exceeding 0.5. When the top predicted allele had at least 50% higher Pattern Score than the second predicted allele (Groups 2 and 4), it was a strong indicator for inducing MMEJ-class repairs. **B** *Top* Definition for each of the 4 groups used in Panel **A**. Each and every zebrafish genomic locus was segmented into these categories. Pattern scores were derived using RGEN online tool. *Bottom* P-values calculated using the Wilcoxon Each Pair Calculation (adjusted for multiple comparisons). **C** Graphical representation of each group detailed in Panel A. Groups 1 and 2 are prone to activate NHEJ-type outcomes, presumably because the yet-unidentified MMEJ factor fails to localize to suitable microhomology arm pairs, limited by how far apart the arms are. Group 4 is most suitable for strong MMEJ activation because it satisfies the proximity requirement AND the relative strength requirement. The latter may aid in the kinetics of the yet-unidentified MMEJ factor binding to the microhomology arms. Our data suggest that Group 3 is an intermediate group in terms of MMEJ activation. Perhaps extragenetic factors, such as cell cycle and epigenetic status may determine how favorable the loci are for MMEJ inductions.

### Accessing the Winner-Take-All Algorithm via MENTHU (MMEJ kNockout Target Heuristic Utility)

The broad potential utility of this updated “Winner-Take-All” Algorithm for MMEJ prediction led us to develop a web-based automated analysis tool called MENTHU (http://genesculpt.org/menthu/). The software can also be downloaded and installed on a local computer (www.github.com/Dobbs-Lab/menthu/). MENTHU accepts a user-specified DNA sequence and targeting scheme as input, and outputs recommended CRISPR sgRNA target sites that are predicted to result in Winner-Take-All type outcomes. We validated the accuracy and functionality of MENTHU against select gRNA sites used in this study using whole exonic sequences as inputs (**S5 Table**). Importantly, the software identified novel Winner-Take-All candidate loci against *surf1* and *tdgf1*, where only Group 3 gRNA loci had been found by previous methods.

## Discussion

To date, precision genome engineering is limited by the ability to predictably, efficiently, and reproducibly induce the identical sequence alterations in each and every cell. Here, we demonstrate the feasibility and utility of creating allelic consistency by an MMEJ-centric approach for designing programmable nucleases. While the precise cellular components of the molecular machinery involved in MMEJ remain incompletely understood(8), we provide evidence that we can enrich for MMEJ events by strictly sequence-based queries.

Importantly, we demonstrate that MMEJ predominant repairs do not operate at the cost of overall mutagenic efficiency; median edit efficiency for Winner-Take-All reagents was 91.4%. As genetically unaltered wild type zebrafish were used throughout the study, we have no reason to believe that NHEJ should have failed at any tested loci. This is in contrast to the current perception that MMEJ is a back-up pathway to NHEJ(7, 8, 16, 17, 27).

Based on the data presented here, we speculate that there is a reaction-limiting factor for MMEJ that is involved in identifying compatible microhomology pairs on both sides of the DNA double stranded break. In the case of abundantly available local microhomology pairs, sometimes this factor fails to localize to a single suitable pair, thus rejecting the MMEJ activation. As end-resection is required for MMEJ and not for NHEJ (9, 17, 18), this yet identified factor may be the deciding factor for committing DSB repair through one End Joining pathway to another. This view is similar to a recent report wherein CtIP/Artemis dependent limited end resection was a key trigger for a slow-kinetic Lig1/3 independent NHEJ event that frequently utilized Microhomology to repair a reporter plasmid(28). In our analysis, the primary driver of this decision making process is the proximity of 2 microhomology arms. Our present findings break down the key triggers for MMEJ activation into a simple 2-component math system with the aims of making the MMEJ-centric approach to gene editing accessible. In addition, this study should inform future studies on the MMEJ pathway by enabling the identification of strongly MMEJ and strongly NHEJ loci under the same genetic context for in-depth comparative molecular analyses.

Successful deployment of the Winner-Take-All reagents makes it possible to directly dictate the reading frame or to do in-frame gene manipulations on endogenous targets. Even assuming a somewhat modest outcome of 50% edit efficiency in which 50% of the mutant allele pool is of the desirable allele, more than 10% of the cell population will be homozygous for this desired allele. Conversely, many real-life gene editing applications would require only one of the diploid copies to be corrected. In these settings under the same assumptions, just 11 viable cells are needed to achieve 95% confidence for establishing the right clone, bringing the idea of this kind of molecular surgery closer to reality.

Our present study expands upon the current state-of-art for MMEJ activation and demonstrates the ability to prospectively design robustly active Winner-Take–All reagents *in-vivo*. We also provide evidence that this 2-component system to identify the Winner-Take-All loci may be broadly applicable beyond zebrafish. To make this algorithm more accessible, we developed the web-based server, MENTHU; http://genesculpt.org/menthu/). MENTHU should enable testing the hypothesis that our present findings are generalizable in other biological systems. In addition, MMEJ-based loci are inherently restricted to genomic locations that leverage endogenous sequence contexts. Consequently, the requirement for a GG dinucleotide for SpCas9 restricts the potential MMEJ sites. However, active investigations are underway to accommodate alternative or more lax PAM requirements. One such example is a recent variant of Cas9 (xCas) that may function efficiently on an NG PAM(29). However, MENTHU allows users to flexibly define any PAM sequence and the cut site (in nts from PAM) to accommodate potential future variants of the CRISPR system.

Finally, we provide strong evidence to support the utility of the MMEJ-centric approach beyond correlating phenotype-genotype in F0 animals. We envision this approach to be useful for: 1) studying the effects of homozygous gene knock-out in culture cells (as opposed to more common, compound heterozygous loss-of-function cell lines), 2) rapid small molecule screening in F0 animals as a complimentary approach to studying in germline mutant animals, 3) globally sharing and reproducing gene knock-out cell and animal lines, and finally, 4) human gene therapy.

## Materials and Methods

### Microhomology arms

For the purpose of this study, microhomology is defined as any endogenous direct sequence repeats of ≥ 3 bp surrounding a DSB site. ≤ 2bp direct sequence repeats were not considered sufficient substrates of MMEJ activation based on our initial analyses of the DSB repair outcomes by previously designed programmable nucleases. Correlation for *Microhomology Fraction* vs the *Slope Value* was tangentially stronger when only ≥ 3 bp arms were considered (r^2^ = 0.382 vs r^2^ = 0.353; **S1 Figure**) in zebrafish, whereas the correlation was lost when 2bp arms were considered in HeLa cells (r^2^ = 0.339 vs r^2^ = 0.034; **S2 Figure**).

### Zebrafish Husbandry

All zebrafish (*Danio rerio*) were maintained in accordance with protocols approved by the Institutional Animal Care and Use Committee at Mayo Clinic. Zebrafish pairwise breeding was set up one day before microinjections and dividers were removed the following morning. Following microinjections, the fertilized eggs were transferred to Petri dishes with E3 media [5mM NaCl, 0.17mM KCL, 0.33mM CaCl, and 0.33mM MgSO_4_ at pH 7.4] and incubated at 28.5°C. All subsequent assays were conducted on fish less than 3 dpf, with the exception of assessing for germline transmission. In this case, injected founders were raised to adulthood per the standard zebrafish husbandry protocol.

### DNA Oligonucleotide Preparation

All of the oligonucleotides used for this study were purchased from IDT (San Jose, CA). Upon arrival, they were reconstituted into 100µM suspensions in 1x TE and stored at −20°C until use.

### sgRNA Expression Vector synthesis

pT7-gRNA was a gift from Wenbiao Chen (Addgene plasmid # 46759). Given that the minimum requirement for the T7 promoter is a single 5’ G, the GG start on this vector was mutagenized to accommodate GA, GC, GT starts, using Forward and Reverse primers given (**S5 Table**). Platinum Pfx DNA Polymerase (Invitrogen 11708013. Carlsbad, CA) was used for 20 cycles of PCR amplification with the Tm of 60 °C and extension time of 3 minutes. DpnI (NEB R0176. Ipswich, MA) was subsequently added to reaction prior to transforming DH5α target sequence was cloned in as previously described, with the exception of cells. The conducting oligo annealing and T4 ligation (NEB M0202. Ipswich, MA) in 2 separate steps. In each case, transformed cells were cultured with Carbenicillin, and plasmids were purified with Plasmid Mini Kit (Qiagen 12123. Hilden, Germany).

### TALEN synthesis

TALEN constructs were generated using the FusX kit (Addgene # 1000000063) as previously described(30). In short, RCIscript-GoldyTALEN was linearized with BsmBI (NEB R0580. Ipswich, MA) along with 6 triplet RVD (Repeat-Variable Diresidue) plasmids. Subsequently, they were ligated together in one reaction by a modified Golden-Gate Assembly. Blue-White colony screening with X-Gal/IPTG, colony PCR and finally pDNA sequencing were done to ascertain the correct assembly.

### In-vitro Transcription and RNA preparation

pT3TS-nCas9n (a gift from Wenbiao Chen: Addgene plasmid # 46757) was linearized with XbaI (NEB R0145. Ipswich, MA), whereas TALEN constructs were linearized with SacI-HF (NEB R3156) and sgRNA vector with BamHI-HF (NEB R3136. Ipswich, MA). Tyr sgRNA #2 – a construct made in the Essner Lab – was linearized with HindIII (NEB R0104. Ipswich, MA). RNA was made using T3 mMessage mMachine kit (Ambion AM1348. Foster City, CA) or HiScribe T7 High Yield RNA synthesis kit (NEB E2040. Ipswich, MA) according to manufacturer’s protocols with the addition of RNA Secure to the reaction (Ambion AM7010. Foster City, CA). To purify RNA, phenol-chloroform extraction was performed using Acid Phenol, Chloroform, and MaXtract High Density Tubes (Qiagen 129046. Hilden, Germany). RNA was then precipitated with Isopropanol at −20 °C, pelleted, air dried and resuspended into nuclease free water. The quality and quantity of RNA were ascertained by using a Nanodrop spectrophotometer and running aliquot on agarose gel. Each batch of RNA was aliquoted into small single use tubes and stored at −80 °C until the morning of microinjections.

### CRISPR-Cas9 RNP preparation for microinjections

sgRNA was thawed on ice in the morning of microinjections. This was then diluted to the concentration of 300 ng/µL in Duplex Buffer [100 mM KCH_3_COO, 30 mM HEPES at pH 7.5]. Appropriate folding of sgRNA was induced by heating it to 95 °C for 5 minutes and gradually cooling the solution to room temperature. Equal volumes of sgRNA and 0.66 mg/mL Alt-R S.p. Cas9 Nuclease 3NLS (IDT 1074181. San Jose, CA) in Cas9 Working Buffer [20 mM HEPES, 100 mM NaCl, 5 mM MgCl_2_, 0.1 mM EDTA at pH 6.5] were mixed and incubated at 37 °C for 10 minutes. RNP solutions were subsequently kept on ice until immediately before use.

### TALEN and CRISPR-Cas9 RNA preparation for microinjections

RNA was thawed on ice in the morning of microinjections. TALEN mRNA was diluted to working concentrations in the range of 12.5 ng/µL to 100 ng/µL in Danieau solution [58 mM NaCl, 0.7mM KCl, 0.4 mM MgSO_4_, 0.6 mM Ca(NO_3_)_2_, 5.0 mM HEPES at pH 7.6]. L L L sgRNA and nCas9n mRNA were mixed and diluted to the final concentrations of 150 ng/µL and 100 ng/µL, respectively, in Danieau solution. These were all kept on wet ice until immediately before use.

### Microinjections

Microinjections were carried out as previously described(31). In short, 1-cell stage fertilized embryos were harvested and aligned on an agarose plate with E3 media. In the case of CRISPR-Cas9 reagents, either 1 or 2 nL was delivered to the cell. In the case of TALEN reagents, 1 ~ 3 nL was delivered to the yolk mass. They were then transferred to Petri dishes in E3 media for incubation at 28.5 °C. Dead and/or nonviable embryos were counted and removed each subsequent morning.

### Phenotype Scoring

Each experiment was conducted in at least a technical triplicate and a biological duplicate. Detailed outcomes are provided in **S4 Table**. Gross phenotypes were scored visually on either 1 dpf or 2 dpf using a standard dissecting microscope. Subsequently, representative pictures were taken with Lightsheet Z.1 (Zeiss 2583000135. Oberkochen, Germany). Shortening Fractions were scored as previously reported. In short, live 2 dpf larvae were immobilized and positioned in 3% methylcellulose. An Amscope camera (MU1403. Irvine, CA) mounted on a Leica Microscope (M165. Wetzlar, Germany) was used to capture a 15 second clip of the beating heart at 66 fps. These clips were subsequently used to measure the distance of the long axis along the ventricle at maximum dilation and maximum contraction using ImageJ software(32). Shortening Fraction was calculated as below:

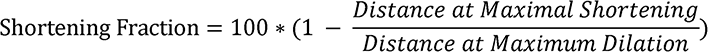

Shortening Fractions from 5 cycles were averaged for each animal.

### DNA extraction and assessing mutagenic outcomes

Typically, 8 uninjected wildtype fish and 8 injected fish were randomly collected without prior screening for phenotype. Chorion was predigested with 1 mg/mL Pronase at room temperature as needed. 1 ~ 3 dpf animals were then sacrificed for individual DNA extractions in 100mM NaOH for 15 minutes at 95 °C. Equal volumes of DNA from the same condition were then mixed and used as templates for PCR with either MyTaq (Bioline BIO-21108. London, UK), Phusion (NEB M0530. Ipswich, MA), or KOD (EMD Millipore 71085. Burlington, MA) polymerases per manufacturer’s protocols. The PCR amplicon was resolved on agarose gel, gel extracted with either Monarch DNA Gel Extraction Kit (NEB T1020. Ipswich, MA) or QiaEx II Gel Extraction Kit (Qiagen 20021. Hilden, Germany), and subsequently sent out for sequencing. The chromatograms from both uninjected and injected amplicons were used for TIDE analysis(22). Alternatively, purified amplicons were used for subcloning analysis with either Topo-TA Cloning Kit (Thermo Fisher Scientific 451641. Waltham, MA) or StrataClone PCR Cloning Kit (Agilent 240205. Santa Clara, CA) per manufacturer’s protocols. Resultant white to pale blue colonies by Blue-White screening were subjected to colony PCR with M13F and R primers, using MyTaq polymerase. Once successful amplification was confirmed on agarose gel, these amplicons were sent out for sequencing either with M13F, M13R or endogenous gene target primers.

### Germline Transmission for 5bp deletion generated by N2B sgRNA #1

RNP containing N2B sgRNA #1 was prepared at 4x diluted dose as described above. Following microinjections, viable fish were raised to sexual maturity. Both F0 founders we attempted to outcross successfully mated and produced viable embryos. DNA was extracted from all viable embryos on 1 dpf, and individual DNA was used as template for PCR amplification using MyTaq Polymerase. Once the thermocycling ran to completion, the amplicons were melted by heating to 95 °C and re-annealed by a gradual step-wise cooling. Surveyor assay(26) was conducted per the manufacturer’s protocol (IDT 706025. San Jose, CA), and the results were analyzed by resolving the post-digest amplicons on agarose gel. Amplicons from 4 heterozygous offspring each were subcloned, and 5 colonies each were sent for Sanger Sequencing to confirm successful transmission of the 5bp deletion allele.

### MENTHU

We developed a software tool, MENTHU (MMEJ kNockout Target Heuristic Utility), to automate calculations required to implement the 2-component Winner-Take-All strategy:1) identification of top predicted microhomology arms separated by ≤ 5 bp of intervening sequence; 2) identification of “low competition” target sites (i.e., with a #1-ranked to #2-ranked Pattern Score ratio ≥ 1.5. We designed MENTHU to first compute two of the same sequence-based parameters (*Pattern Score and Microhomology Score*) used in the algorithm of Bae *et al*., (which are computed online by the RGEN online tool, http://www.rgenome.net), by re-implementing and modifying the original Python source code provided in Supplemental Figure 3 (14) in R. The MENTHU webserver operates under R(33) version 3.4.1 and RShiny (34) v1.0.5. The MENTHU code was built through RStudio(35) v1.1.442. Details regarding specific R package versions, complete documentation and a full downloadable version of MENTHU for local installation are provided at www.github.com/Dobbs-Lab/menthu/. MENTHU v2.0 can be freely accessed online at http://genesculpt.org/menthu/.

### Statistical Analyses

All of the statistical analyses were carried out using JMP software (SAS Institute. Cary, NC). In all instances, p-values were calculated assuming non-Gaussian Distributions. Wilcoxon Each Pair calculation was used for multiple group comparisons with adjusted p-values.

## Acknowledgements

Funding: NIH OD020166; NIH UL1TR002377; NIH GM63904; P30DK090728;

P30DK84567; AHA 16PRE30470004; Mayo MSTP; Mayo Foundation; and gift from Marriott Foundation.

We would like to acknowledge Melissa McNulty for TALEN synthesis and data curation, Mark Urban for zebrafish husbandry and help with microinjections, Patrick Blackburn for designing *chrd* #1 sgRNA, Mayo Clinic Zebrafish Facility and staff for their support. We also thank the Jin-Soo Kim group for developing the Microhomology-Predictor CRISPR RGEN Tool, for making source code freely available, and for sharing the deep sequencing output from their HeLa cell experiments. We thank Wesley A Wierson and Jeffrey J Essner for the tyr gRNA, and the research groups of Dr. Essner and Dr Maura McGrail for valuable discussions and feedback and for MENTHU server testing, and Carolyn Lawrence-Dill and her group, especially Darwin Campbell, for valuable discussions and hosting services for MENTHU.

---

**S1 Figure Overabundance of Microhomology arms is a negative predictor of MMEJ activation in zebrafish**

**A** Box plot showing the distribution of *Slope Values* across 19 zebrafish genomic targets. **B** Scatter plot of 3bp MH Fraction against *Slope Value*. Linear fit with 95% Confidence Interval (shade) is shown. r^2^ = 0.382, p = 0.0048. **C** Scatter plot of 2bp MH Fraction against Slope Value. Linear fit with 95% Confidence Interval (shade) is shown. r^2^ = 0.353, p = 0.0073 Shade is 95% CI. *Pattern Scores* and *Microhomology Scores* were derived using RGEN online tool (http://www.rgenome.net).

**S2 Figure Overabundance of Microhomology arms is a negative preidictor of MMEJ activation in HeLa cell**

**A** Scatter plot of 3bp MH Fraction against *Slope Value* from the first 50, alphabetically sorted HeLa cell targets. Linear fit with 95% Confidence Interval (shade) is shown. r^2^ = 0.339, p = 0.0001. **B** Scatter plot of 2bp MH Fraction against *Slope Value* from the first 50, alphabetically sorted HeLa cell targets. Linear fit with 95% Confidence Interval (shade) is shown. r^2^ = 0.034, p = 0.2644. **C** Box plot showing the distribution of Slope Values across the first, alphabetically sorted HeLa cell targets. **D** Box plot showing the 3bp MH Fractions for High and Low competition sites amongst the remaining 40 HeLa cell targets. p = 0.011. Targets with < 20% overall edit efficiency were excluded in all panels. *Pattern Scores* and *Microhomology Scores* were derived using RGEN online tool (http://www.rgenome.net).

**S3 Figure Microhomology allele generated by *ttn.2* N2B sgRNA #1 is germline transmitted**

Agarose gel showing PCR amplicon post Surveyor digest. 752bp band is the whole amplicon. The expected bands due to mutations at the CRISPR site are denoted by yellow arrowheads. The red asterisk denotes positive digest band due to a background T -> A SNP at position 389 from the 5’ end of the amplicon. Heterozygous animals are bolded and underlined. Genotypes of the first 4 heterozygous progenies from each founder were ascertained by subcloning analyses.

**S4 Fig Fitting Competition Hypothesis Version using HeLa cell dataset**

Outlier plot summarizing repair outcomes from 90 genomic targets using CRISPR-Cas9. Similar to the findings in zebrafish, close proximity of 2 MH arms (Groups 3 and 4) appears to be the primary determinant for utilizing this MH pair efficiently. When the top predicted allele had at least 50% higher *Pattern Score* than the second predicted allele (Groups 2 and 4), median Top MH Fractions trended higher compared to Group 1 and 3, respectively. P-values calculated using the Wilcoxon Each Pair Calculation (adjusted for multiple comparisons). Targets with < 20% overall edit efficiency were excluded from analysis. *Pattern Scores* and *Microhomology Scores* were derived using RGEN online tool (http://www.rgenome.net).

**S1 Table**

List and summary mutagenic outcomes of TALEN and CRISPR-Cas9 reagents that were designed primarily using the Bae *et al*. algorithm(14). Underlined & italicized bases in sgRNA sequence denote mismatched bases due to the promoter requirement.

*Pattern Scores* and *Microhomology Scores* were derived using RGEN online tool (http://www.rgenome.net).

MH: Microhomology, SC: Subcloning

^*^Reagents prospectively designed according to Bae *et al*. algorithm(14).

^†^No raw sequencing data were available. However, the outcome had been compiled into a table prior to conception of this study.

^‡^Injected with sgRNA and Cas9 mRNA (150 pg and 100 pg, respectively)

^^^gift from Wenbiao Chen (Addene # 46761).

**S2 Table**

Summary gross phenotyping outcomes from Winner-Take-All reagent injections. For *tdgf1*, Experiments 1a and 1b correspond to technical replicates using WT 1 as reference, uninjected control. *chrd* and *tdgf1* phenotypes were scored on 1 dpf, whereas *tyr, ttn.2* N2B, *ttn.2* phenotypes were scored on 2 dpf.

**S3 Table**

List and summary sequence outcomes of Low Competition sgRNA that were designed around the Competition Hypothesis. Underlined & Italicized bases in gRNA sequence denote mismatched bases due to the promoter requirement. *Pattern Scores* and *Microhomology Scores* were derived using RGEN online tool (http://www.rgenome.net). MH: Microhomology, SC: Subcloning, TIDE: Tracking Indels by DEcomposition.

^†^ injected RNP at the dose of 115 pg sgRNA and 245 pg Cas9 due to poor viability at higher doses

**S4 Table**

List and summary sequence outcomes of Medium ~ High Competition sgRNA that were designed around the Competition Hypothesis. Underlined & Italicized bases in sgRNA sequence denote mismatched bases due to the promoter requirement. *Pattern Scores* and *Microhomology Scores* were derived using RGEN online tool (http://www.rgenome.net).

MH: Microhomology, SC: Subcloning, TIDE: Tracking Indels by Decomposition.

^†^injected RNP at the dose of *115 pg sgRNA and 245 pg Cas9* due to poor viability at higher doses

**S5 Table**

Sample MENTHU output from select CRISPR-Cas9 targets used in this study. The output was obtained by using the entire target exon sequence with 40bp intronic sequence each on both 5’ and 3’ ends. The MENTHU output provides a 3’ NGG PAM sequence for each gRNA targets (italicized and underlined). MENTHU gRNA outputs that matched the target sequences used in this study are bolded. *Criteria 1* and *2* refer to 1) if top predicted microhomology arm is separated by 5bp or less, and 2) if the ratio of top to second predicted pattern scores is at least 1.5. MENTHU is programmed to terminate calculations if the target site is negative for Criterion 1. As a result, no gRNA sequence output is obtained for *chrd* #1 and *mitfa* #2. Importantly, in two instances (*surf1* and *tgdf1*) where we only had Group 3 reagents, novel candidate Winner-Take-All sites were identified.

**S6 Table**

List of primers used in this study. All the primer sequences are provided in 5’ → 3’ order. For *urod* Reverse primer, M13F primer sequence was added at the 5’ end of the endogenous target sequence (bolded and italicized). For SDM primers, intended point mutation is indicated by bold and italic.

